# The Role of Metabolic Strategies in Determining Microbial Community Diversity along Temperature Gradients

**DOI:** 10.1101/2024.08.28.610078

**Authors:** Quqiming Duan, Tom Clegg, Thomas P. Smith, Thomas Bell, Samraat Pawar

**Affiliations:** Imperial College London; Helmholtz Institute for Functional Marine Biodiversity at the University of Oldenburg (HIFMB)

## Abstract

By conservative estimates, microbes make up about 17% of the world’s biomass and are essential for most ecosystem functions. However, the mechanisms driving the variation in microbial species diversity in response to both natural and anthropogenic temperature gradients remain unclear. In this study, we integrate ecological metabolic theory with a community assembly model to predict that microbial community diversity generally follows a unimodal pattern with temperature. The position and magnitude of peak diversity are determined by interaction-driven species sorting acting on variations in the temperature dependence of carbon use efficiency (CUE) and generalist-specialist tradeoff. Specifically, trait sorting across temperatures favours communities with high mean and low variance in species-level CUEs. We provide empirical evidence supporting our qualitative predictions of the unimodal temperature-diversity pattern along the global geological temperature gradient, which peaks at about 10-15 ^*°*^C. Our findings indicate that the response of diversity as well as CUE to temperature of microbial communities can be predicted from relatively feasible life-history trait measurements, paving the way for interlinking microbial community diversity and carbon cycling along spatial and temporal thermal gradients.

## 1 Introduction

Microbes, encompassing bacteria, archaea, fungi, protists, and viruses, are highly diverse (Locey & Lennon 2016) and ubiquitous in our biosphere. Among these, bacteria dominate, making up approximately 13% of global biomass (Bar-On et al. 2018), accounting for a variety of ecosystem functions by metabolising substrates in their environment (Handley 2019), and ultimately driving all major global biogeochemical cycles, including carbon (Bardgett et al. 2008) and nitrogen (Stein & Klotz 2016). The taxonomic diversity of the microbial communities encodes the diversity of functional traits and therefore plays a critical role in determining their functional ability (Wagg et al. 2019, Graham et al. 2016, Fierer 2017, Bell et al. 2005). However, the emergence and maintenance of biodiversity in microbial communities under varying natural and anthropogenic environmental conditions remains poorly understood.

All organisms obtain energy and matter for their growth and reproduction through cellular enzyme-catalysed metabolic reactions. Because temperature is the primary driver of all chemical reactions, it also plays a fundamental role in determining metabolic traits at the species level, interactions between species, and ultimately their fitness in a community context (Arroyo et al. 2022, Dell et al. 2011, Rezende & Bozinovic 2019, Smith et al. 2021). Indeed, environmental temperature on Earth is highly variable on both geographic (e.g. latitudinal, altitudinal) and temporal scales (e.g. daily to seasonal cycles, anthropogenic climate change) and is known to be a major determinant of both contemporary and historical biodiversity patterns across taxonomic groups and habitats (Kreft & Jetz 2007, Yasuhara & Danovaro 2016, Antão et al. 2020, Mayhew et al. 2008, 2012). Yet, the mechanistic link between environmental temperature and microbial community diversity remains largely unknown (Zhou et al. 2013, Spicer et al. 2019, Helmuth et al. 2005). Furthermore, previous empirical studies have reported conflicting results on the shape of the microbial temperature-diversity pattern (Hendershot et al. 2017), finding increasing (Zhou et al. 2016), decreasing (Kolton et al. 2019), unimodal (Singh et al. 2012, Milici et al. 2016) and temperature insensitive responses (Zhou et al. 2020, Fierer & Jackson 2006).

In this study, we integrate metabolic theory of ecology with a community assembly model to predict microbial community diversity, and investigate the mechanisms underlying the temperature-diversity pattern. In particular, we focus on species-level carbon use efficiency (CUE)(Sinsabaugh et al. 2013, Manzoni et al. 2012) as a fitness-defining trait to study community assembly along temperature gradients. CUE quantifies the efficiency of resource utilisation, which is fundamental in determining species’ population growth and relative fitness within a community and is highly sensitive to changes in environmental temperatures (Smith et al. 2021, Pold et al. 2020).

## 2 Results

### Model

We model the temperature dependent dynamics of microbial communities using a modified version of the Microbial Consumer-Resource Model (MiCRM) (Fig 1) (MacArthur 1970, Marsland III et al. 2019, Lechón-Alonso et al. 2021, Cui et al. 2020). The MiCRM modelles the resource-mediated dynamics of microbial consumers and is able to capture the complex interactions that arise between populations through both competition for resources and the exchange of metabolic by-products.

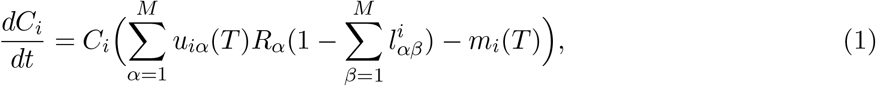

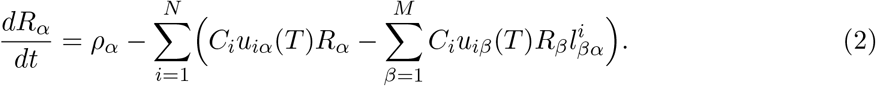

Here, *C*_*i*_ and *R*_*α*_ are the biomass and abundance of the *i*th consumer and *the α*th resource in the environment, respectively. Each consumer uptakes carbon resources from the surrounding environment according to its temperature-dependent (*T*) metabolic preference *u*_*iα*_(*T*). A proportion of this uptake is used for biomass accumulation and the rest is leaked due to inefficiency or as metabolic by-products (whose identity is determined by the tensor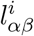. Consumers also experience an additional temperature-dependent loss term *m*_*i*_(*T*) that represents maintenance costs (particularly through respiration). Resources are supplied at some rate *ρ* and then taken up and released as by-products by consumers as described above.

**Figure 1.**
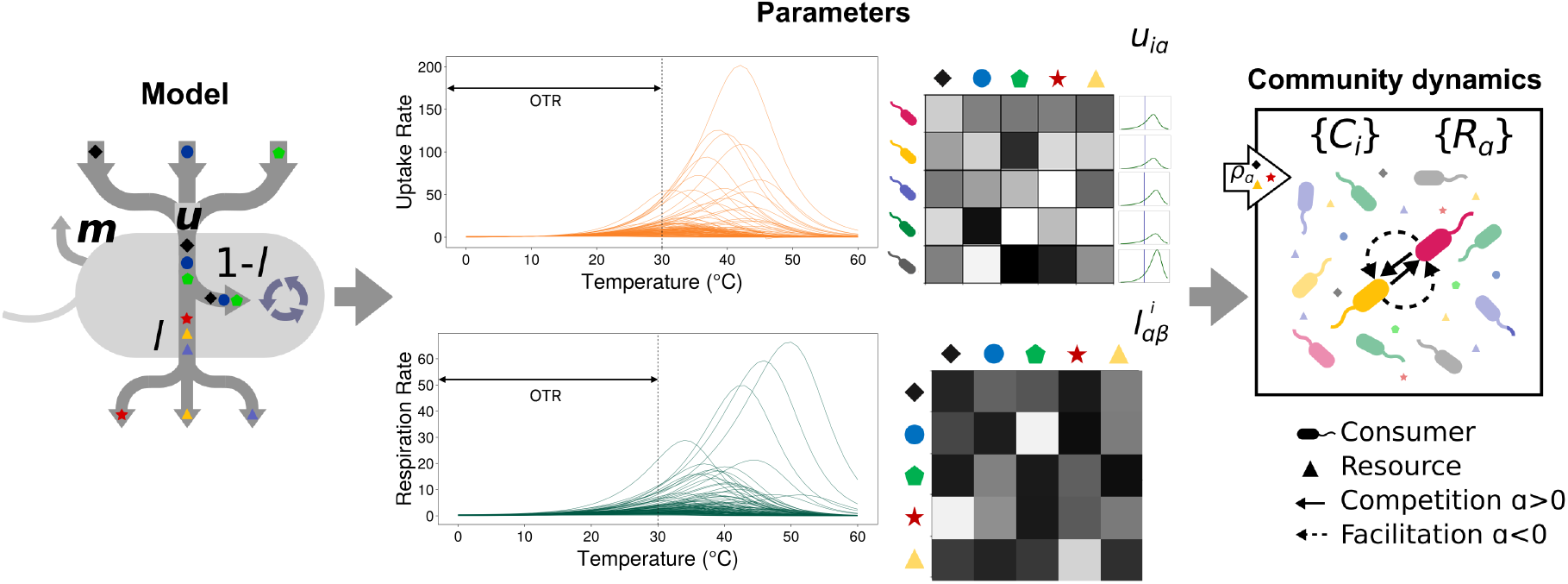
Schematic of the Microbial Consumer-Resource Model. The dynamics of consumer abundances (*C*_*i*_s) are determined by carbon gain through resource uptake (with the rate *u*), and loss through leakage/transformation (*l*) and maintenance respiration (with the rate *m*). These species’ level resource uptake rates and maintenance respiration rates are constrained by unimodal temperature performance curves (TPCs). The black arrows in TPC plots suggest species’ operational temperature ranges (OTR), which is the temperature range species normally experience in their life cycles before high-temperature effects come into play for most species. Resource abundances (*R*_*α*_) are constantly supplied by the inflow of external resources supply (*ρ*). Species within the community compete through the exploitation of resources and cross-feed through the secretion of metabolic by-products, which replenish the resource pool.

We model the temperature dependence of consumer uptake preference *u*_*iα*_(*T*) and maintenance costs *m*_*i*_(*T*) using the modified Sharpe-Schoolfield equation (see Methods). This describes the unimodal thermal response of biological traits in which rates tend to increase exponentially with rising temperature up to a limit after rates fall since enzymes for biochemical reactions deactivate at high temperatures. By including this temperature dependence, we not only introduce a direct effect of temperature on consumers but also allow an indirect effect to emerge through the interactions between populations. As we will show, this in turn scales up to affect community-level emergent properties such as richness we focus on here.

### Microbial Thermal Strategies and Species Sorting

Species’ carbon use efficiency (CUE) is essentially 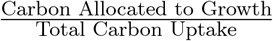, and includes the partitioning of carbon resources harvested into cell growth versus maintenance loss (Geyer et al. 2016) (Equation 4). CUE is the intrinsic property of a given species governed by its cellular metabolic network, and reflects a major component of its life history strategy to maximise fitness (henceforth, “metabolic strategy”) (Manzoni et al. 2012). The temperature dependence of CUE is a function of the temperature dependence of the underlying metabolic traits (Smith et al. 2021, Qiao et al. 2019). Species in natural environments typically operate within a temperature range (the operational temperature range, OTR) that is typically a few degrees lower than their optimal growth temperature (Smith et al. 2019). Within this OTR, species uptake and respiration rates increase exponentially with increasing temperature. Therefore, for most species in any given local environment, the CUE of species also increases exponentially with temperature.

At each temperature along a temperature gradient, species with higher resource uptake rates and lower respiration rates (higher CUE) typically succeed during species sorting (Figure 2 a-b). This can be explained by the competitive exclusion principle (the R* rule, Tilman 1982) because species with the highest CUE can deplete resources to a lower resource level to grow and survive (Figure 2 c, Equation 6). Thus, in general, at any given temperature along a gradient, species sorting inevitably results in a subset of species from the local or global pool with the highest CUE in that thermal environment.

**Figure 2.**
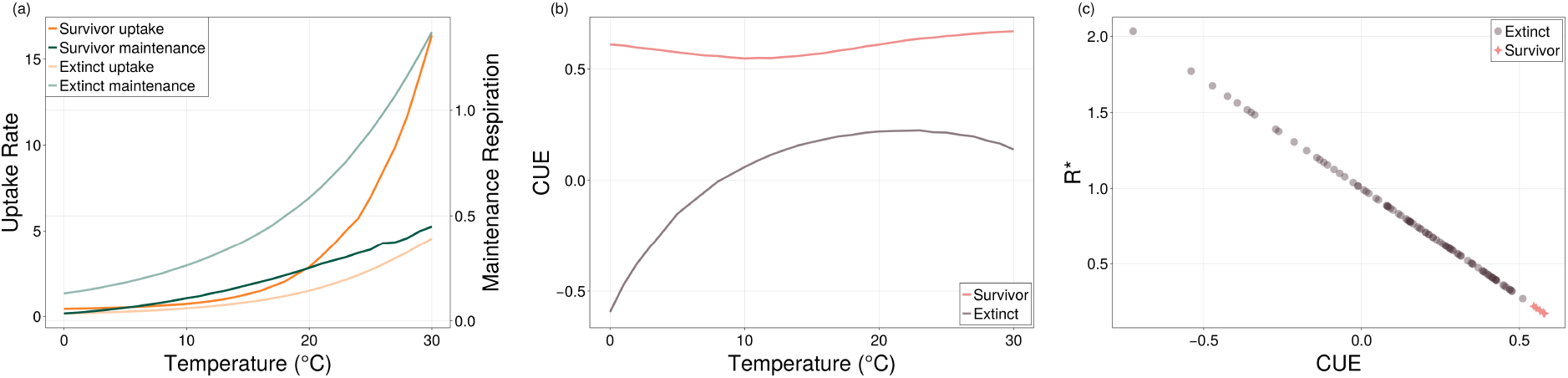
Temperature-dependent species sorting selects for specific life history traits along thermal gradients. (a) Species with relatively higher resource uptake rates (orange lines) and relatively lower maintenance respiration rates (green lines) typically survive. (b) Species with the highest CUE survive at all temperatures. The red line depicts the mean CUE of survivors, and the grey line depicts the mean CUE of species excluded in species sorting. (c) The negative correlation between species’ CUE and their equilibrium resource requirements (*R*^∗^) at an example temperature (15 ^*o*^C). The red dots depict species that survived competitive exclusion, and the grey dots represent excluded species. All mean values in this figure are taken from 670 simulations at each temperature, plotted with standard error.

### Local and Global Temperature-Diversity Patterns

The generation time of most bacterial species in natural environments ranges from minutes to months (Gibson et al. 2018). Therefore, species typically experience various subsets of the constantly fluctuating environmental temperature. Species adapted to significant thermal fluctuations across generations tend to evolve into thermal generalists (Kontopoulos, Smith, Barraclough & Pawar 2020), with relatively lower activation energies (*E*, the rate at which species metabolic traits increase with temperature) and relatively higher low-temperature performances (*B*_0_) (Gilchrist 1995). There is almost no correlation between *E* and *B*_0_ (correlation coefficient *ρ* ≈ 0) for these species. On the other hand, for significant thermal fluctuations within the generation or for constant environments, thermal specialists are favoured (Kontopoulos, Smith, Barraclough & Pawar 2020, Gilchrist 1995), with relatively higher *E* values and relatively lower *B*_0_ values. This results in a strong trade-off between *E* and *B*_0_ (correlation coefficient *ρ* ≈ 1) within species’ TPCs. Here, we consider both extreme correlation types when investigating the deterministic role of species’ thermal performances of metabolic traits on local microbial diversity-temperature patterns.

Figure 3 shows the two extreme cases of how these covariances in species’ TPCs affect microbial diversity along temperature gradients on the geological scale. Figure 3a shows a scenario in which species are from warmer and less fluctuating temperatures (e.g., closer to tropic regions on natural latitudinal gradients) (Pawar 2005, Kontopoulos, Smith, Barraclough & Pawar 2020, Kontopoulos, van Sebille, Lange, Yvon-Durocher, Barraclough & Pawar 2020). These communities are mainly dominated by thermal specialists who have narrower temperature breaths with higher fitness at their preferred temperatures (Huey & Hertz 1984, Angilletta 2009, Deutsch et al. 2008). These species typically have smaller thermal safety margins, and their preferred temperatures are relatively closer to their respective optimal temperatures. A strong trade-off between *E* and *B*_0_ is typically seen in their thermal performance curves (*ρ* ≈ 1). In this case, variance of species’ life history traits is minimised at a relatively high temperature, permitting a larger subset of species to coexist there. However, for communities from cooler (e.g., temperate) areas (scenario in Fig. 3b), a steep thermal performance curve is not as advantageous (MacLean et al. 2019). Thermal generalists, who perform relatively equally across all temperatures, can also achieve the minimum fitness requirement for survival at these relatively cooler temperatures. This results in a relatively higher variation in the traits of the species in general. Greater intersections of species’ metabolic trait TPCs close to the centre of their OTR are expected. At these intersections, the species richness also peaks because it is there that the difference in species’ relative fitness within the community is minimised.

**Figure 3.**
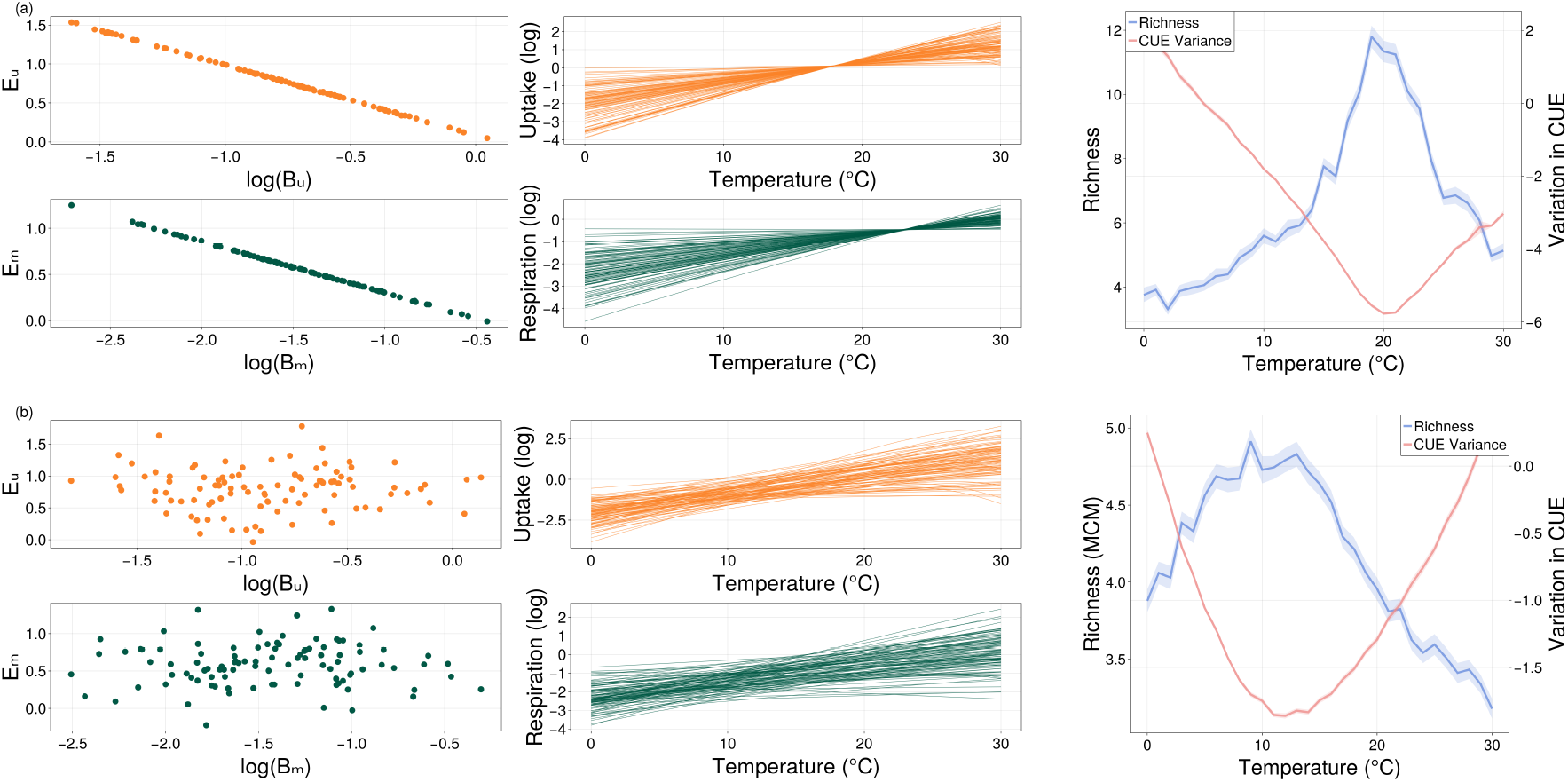
Difference in species’ thermal strategies (i.e., species’ CUE) generate different diversity patterns with temperature along temperature gradients. The variance in CUE along the temperature gradient was changed by varying the covariance between species’ *B*_0_ (the trait value at the relatively low reference temperature) and *E* (the activation energy). Here, we show examples with correlation coefficients *ρ* as -1.0 (a), 0.0 (b). (a) For communities residing in relatively hotter environments, the covariance between *B*_0_ and *E* is strongly negative, and the variation in species’ life history traits is lowest at relatively higher temperatures where richness peaks. (b) For communities residing in medium to cooler natural environments, the covariance between species’ *B*_0_ and *E* is almost zero. Richness shows a relatively weaker correlation with changing temperature, and the variation of species traits is at the lowest at a relatively cooler temperature where richness peaks.

We propose that the overall response of diversity of microbial communities to a temperature gradient over space, given sufficient time for species sorting and adaptation to local conditions (such as the mean or median environmental temperature, the amplitude and frequency of temperature fluctuation within that region), will ultimately follow a unimodal pattern when measured over a sufficient temperature range. The temperature at which this diversity curve peaks is determined by the distribution of the TPCs of the traits that underlie CUE. More specifically, it is determined by the correlation between *B*_0_ and *E* of these traits, where a negative covariance is expected under the generalist-specialist trade-off.

Using a global database of trait TPC (Smith et al. 2019, 2021), we predict a unimodal pattern of richness along the global geological temperature gradient that peaks at a relatively low temperature of approximately 10-15 ^*o*^C (Figure 4). This prediction aligns well with the temperature-diversity pattern recorded by the Earth Microbiome Project (Thompson et al. 2017).

**Figure 4.**
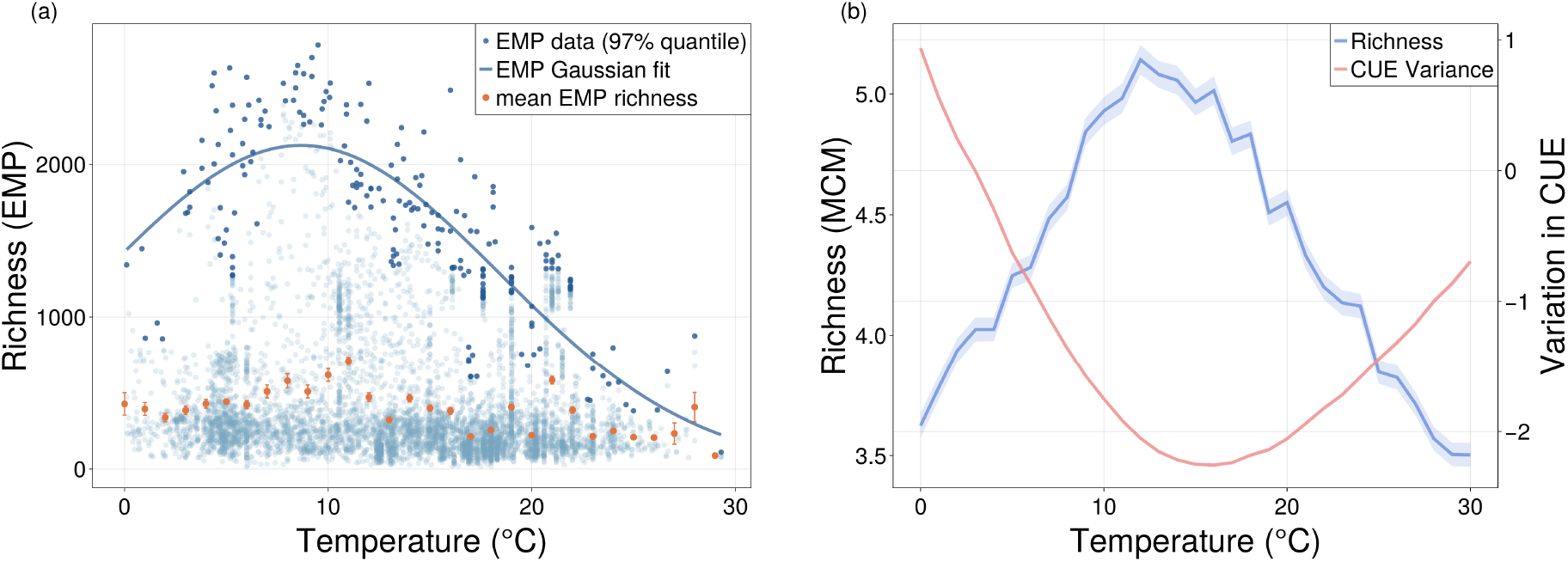
The unimodal pattern of microbial diversity along the temperature gradient. (a) The temperature-diversity pattern predicted by the Earth Microbiome Project (EMP). Each blue dot represents the richness of an empirically collected community; dark blue dots are the 97% quantile of this biodiversity data. The blue line represents a fit of the Gaussian equation to the 97% quantile data, and the orange dots track the mean and standard error of species richness along this subset temperature gradient of 0 to 30 ^*o*^C. (b) The variation of species CUE drives the unimodal pattern in microbial diversity along the temperature gradient. The blue line depicts the unimodal temperature-diversity pattern, and the red line is the mean log variance of species CUE within the initial community before species sorting. Both lines are plotted with mean values of 670 simulations at each temperature with standard error.

## 3 Discussion

We have investigated the temperature richness pattern using a novel, mechanistic microbial community assembly model parameterised with empirically constrained thermal dependencies of bacterial carbon use efficiency and its underlying metabolic traits. The results show that, in general, microbial community richness, when measured across a sufficiently wide thermal gradient, will follow a unimodal pattern with temperature. This pattern, along with the average community CUE, is always predictable if given the TPCs of life-history traits (e.g. growth rate, resource uptake rate, respiration rate) at any empirical site. Our parameterisation using the global dataset of bacterial traits indicates that, in general, species richness globally is expected to peak at a relatively cool temperature between 10-15^*o*^C.

Our study has general implications for understanding biodiversity under both long-term (e.g., anthropogenic global warming) and short-term temperature fluctuations and the emergent geological distributions of global bacterial diversity. For example, in our prediction of the pattern of microbial diversity along global temperature gradients (Figure 4), an increase in temperature, such as annual warming caused by ongoing climate change, could induce diversity loss, especially in temperate and tropical regions. In all thermal regimes, species sorting is vital in determining the emergence and response of microbial communities to temperature fluctuations. Species that are outcompeted at certain temperatures could still exist within the species pool as “latent functional diversity” (Smith et al. 2022), dormant and waiting for their preferred (or preadapted) thermal environments, where species sorting would once again favour these species. Our model predicts that the pattern of microbial diversity along the global temperature gradient is always unimodel within when measured over a sufficiently wide temperature range. The ostensibly conflicting findings on the microbial temperature-diversity pattern reported by previous empirical studies (Hendershot et al. 2017) are likely due to the varying temperature ranges examined in the regions studied, which would result in different subsets of the unimodal temperature diversity curve. Consistent with our prediction, microbial community diversity reported by the Earth Microbiome Project (EMP)’s global survey (Thompson et al. 2017) exhibits a unimodal pattern that peaks at relatively cool temperatures (Figure 4). Thompson et al. (2017) also reported nestedness wherein the composition of communities with fewer species is generally a subset of their more diverse counterparts, suggesting that species sorting plays an important role in driving microbial diversity. It is important to note that our prediction of microbial diversity along the global geological temperature gradient is a general macroecological pattern. It is a compilation of all temperature-driven microbial diversity patterns of local communities along the entire gradient. These local communities are the result of species sorting based on species’ thermal metabolic strategies in the respective species pool. Additionally, we have validated our prediction of this global geological temperature-diversity pattern by incorporating the evenness of species abundances, using the Shannon and Simpson diversity indices (Figure SI.2).

The temperature for the peak diversity that we predict coincides with the peak diversity temperature in the mean richness values along global temperature gradients, but slightly deviates from the maximum richness temperature observed in the 97% quantile of the EMP data. This probably stems from i) biases in our global dataset of TPCs, which do not contain the TPCs of resource uptake rates, even though these theoretically align with the growth rates of species, and the specific measurements of resource uptake rates might result in small differences in the variation and the covariance between normalisation constant and activation energy, shifting the diversity peak along the temperature gradient. ii) Biases in the EMP sampling, such as the relatively extensive sampling in temperate regions. iii) Other environmental factors that could have an effect on the diversity of the local community, such as climate (e.g., precipitation), geochemical and resource contents at sampling sites, sampling seasons, and habitat types of focus.

We stress the importance of bringing together population dynamics and metabolic theory of ecology (Brown et al. 2004), that community-level properties such as diversity are species sorting results based on species’ metabolic strategies, rather than a summation of species-level temperature dependencies as previously proposed (Arroyo et al. 2022). Previous theoretical studies that attempted to bring these together in predicting species diversity along temperature gradients used a phenomenological model, the generalised Lotka-Volterra model (GLV). They incorporated the temperature dependencies of metabolic rates into this model (Stegen et al. 2012, Clegg & Pawar 2022), which also provided unimodal patterns of temperature diversity. With an effective version of our MiCRM, we were able to produce a similar pattern predicted by Clegg & Pawar 2022, who calculated the probability of feasibility along temperature gradients using a mean field approximation (Figure SI.3). However, the “unrealistic” GLV model (Levins 1966) purely incorporated the observed competitive interactions between species, without explaining the underlying mechanism of these interactions in terms of metabolic strategies. Another study using a similar version of the consumer-resource model also demonstrated a unimodal pattern of temperature-diversity pattern (Marsland et al. 2020), but only reproduced the pattern observed in EMP by simply increasing the maintenance costs of all species for temperatures away from the optimal temperature recorded in Thompson et al. (2017), without discussing the mechanisms that determined this biodiversity pattern.

The value of diversity and richness in our global prediction (Figure 4) is generally lower compared to the empirical field measurements for the following reasons. Firstly, we started from a system of just 100 species competing for 50 types of resource, which is sufficient to generate qualitatively robust predictions. In addition, our species pool is not classified by OTU, but by functional groups, where in general the bacterial community assemblies converge on functional groups based on metabolic niches (Pascual-García et al. 2020, Goldford et al. 2018). Therefore, our richness measurements are more representative of functional diversity, where each functional group contains various taxonomic-level species (Louca et al. 2016, Burke et al. 2011).

We ran our assembly simulations only within 0-30^*o*^C because the OTR of most ectotherm species lies within this range (Corkrey et al. 2016, Smith et al. 2019), and this temperature range covers most of the global annual temperatures (Karger et al. 2017). Our predictions can be extended to higher temperatures, provided that data sets of trait TPCs are available, such as those relevant to complex microbiomes that reside within homeotherms (e.g., the human microbiome Lloyd-Price et al. 2017) that normally operate in a higher temperature range of approximately 36 - 42 ^*o*^C (Ivanov 2006) and species that reside in hot springs that are adapted to temperatures up to boiling (Podar et al. 2020). Extending our prediction towards lower temperatures (e.g., the polar regions) would require the consideration of low-temperature deactivation (Schoolfield et al. 1981, DeLong et al. 2017) within the temperature performance function.

As a result of species sorting, we suggest that there is no universal advantage in any particular shape of the CUE-temperature pattern along temperature gradients (Pold et al. 2020), while species with relatively the highest CUE value always outcompete at any given temperature. Therefore, for the same species pool, thermal generalists might have a better chance of succeeding at relatively lower temperatures, and increasing temperatures would result in a community mainly composed of thermal specialists. We have shown that with the constant exclusion of species with relatively lower CUE at all temperatures, the average species-level CUE within the community does not respond significantly to changing temperatures. This constant ecological selection in species with relatively high CUE is possibly due to a general lack of abiotic stress and resource limitation in our simulated environment (Malik et al. 2020). For communities living in stressful environments, such as chemical or physical stresses (e.g., draught) or a deficiency of simple degradable resources, species tend to partition energy into stress tolerance (maintenance costs) or resource acquization, instead of increasing biomass yields (Schimel et al. 2007). In these environments, species with higher tolerance or specific biodegradability would be favoured.

In conclusion, our findings indicate that community assembly along temperature gradients is driven by species’ life-history strategies, particularly their metabolic traits and CUE. Coexisting species at each temperature are those with the highest CUEs that are thermodynamically and metabolically feasible at that temperature. Our study offers a relatively simple framework for mechanically linking microbial diversity and functioning with local to global carbon cycling. For future empirical studies on temperature effects on community dynamics (e.g., climate change, etc.), our work highlights the importance of measuring the thermal performance curves of a representative set of species’ metabolic traits. This approach would allow an adequate charting of the distribution of species’ thermal strategies within a particular community. In addition, we expect the rapid development of methodologies and technologies in empirical studies to validate and refine our theoretical understanding and prediction.

## 4 Methods

### 4.1 The Microbial Community Model

We perform microbial community assemblies using the Microbial Community Model (MiCRM), a McArthur-style consumer-resource model (MacArthur 1970, Marsland III et al. 2019, Lechón-Alonso et al. 2021, Cui et al. 2020) through random immigration of microbes from a global species pool with different thermal physiologies of metabolic strategies along the temperature gradient. This model tracks carbon fluxes in a consumer-resource system with metabolic mechanisms such as resource uptake, metabolic by-product secretion (including facilitation mechanisms such as cross-feeding among species), and maintenance respiration. For *N* species competing for *M* resource types, the biomass dynamic of species *i* and the resource dynamic of resource type *α* is given by:

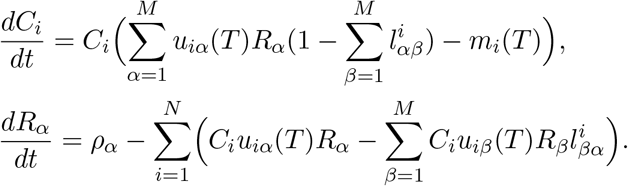

Here, *C*_*i*_ denotes the biomass abundance of the *i*^*th*^ consumer and *R*_*α*_ denotes the resource abundance of the *α*^*th*^ type of resource. In this set of equations, species growth is determined by the uptake of preferable resources (*u*_*i*_) and maintenance loss mainly through respiration (*m*_*i*_). The uptake preferences of each consumer are encoded in an uptake preference matrix, in which *u*_*iα*_ denotes the uptake preference of the resource *α* by species *i*. For resource uptake at the species level, a proportion of resource abundance consumed by each species (for example, species *i*) is lost through inefficiency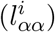 or transformation into metabolic by-products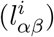 that is then returned to the resource pool and become available carbon sources for other species. Constraints on assimilation efficiency are set for both resources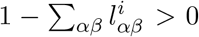 and species 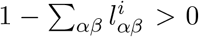. The resource abundance dynamics are controlled by the input through the constant external supply of all types of resources plus the secretion of metabolic by-products; and the output, which is species uptake of carbon resources according to their preference matrix *u*_*iα*_. The parameter descriptions are given in table 1.

**Table 1:** Definitions for parameters in the microbial community model.

| Symbols | Definition |
| --- | --- |
| $M$ | Number of resource types |
| $N$ | Number of consumer species |
| $C_i$ | Biomass concentration of consumer $i$ |
| $R_{\alpha}$ | Resource concentration of resource $\alpha$ |
| $u_{i\alpha}$ | Uptake rate of consumer $i$ on resource $\alpha$ |
| $m_i$ | Maintenance loss of consumer $i$ |
| $l_{\alpha\beta}^i$ | Leakage/transformation fraction of resource $\alpha$ to $\beta$ by species $i$ |
| $\rho_{\alpha}$ | External resource supply of resource $\alpha$ |

In these equations, the resource uptake rate and maintenance respiration rate for each species are temperature-dependent and are denoted by (*T*). The equations for their temperature dependencies are shown in the following section.

### 4.2 Temperature Dependency of Metabolic Traits

Species metabolic rates typically display unimodal shapes with temperature (Kontopoulos, van Sebille, Lange, Yvon-Durocher, Barraclough & Pawar 2020, Angilletta 2009), which increase exponentially with rising temperatures toward their thermal optimum followed by an exponential decrease due to high-temperature effects. These are normally depicted as temperature performance curves (TPCs). We describe the TPCs of metabolic rates (resource uptake rate and maintenance respiration rate) using the modified Sharpe-Schoolfield equation (Kontopoulos, van Sebille, Lange, Yvon-Durocher, Barraclough & Pawar 2020):

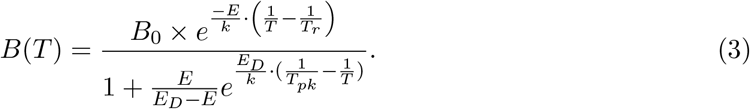

Here, *B*(*T*) denotes the trait value at a given temperature *T* and *B*_0_ denotes the normalisation constant, the value of which is the metabolic rate value at reference temperature *T*_*r*_. As the temperature increases, the metabolic trait increases exponentially following the rate *E* (which is the activation energy of typically responsible enzyme reactions) until their optimum performance temperature, then the acceleration drops as enzymes begin to deactivate at higher temperatures and the trait value reaches its highest performance at *T*_*pk*_. As the temperature continues to increase and exceeds *T*_*pk*_, the value of metabolic traits decreases exponentially following the rate *E*_*D*_, which is the apparent deactivation energy. The parameter descriptions are given in table 2.

**Table 2:** Parameters for the temperature dependencies of uptake and respiration.

| Symbols | Definition | Units |
| --- | --- | --- |
| $B_0$ | Normalization constant | - |
| $E$ | Activation energy | eV |
| $k$ | Boltzmann constant | eV/K |
| $T$ | Model temperature | K |
| $T_r$ | Reference temperature | K |
| $E_D$ | Deactivation energy | eV |
| $T_{pk}$ | Peak temperature for metabolic rates | K |

The Sharpe-Schoolfield equation is chosen here to approximate more realistic metabolic rates at all temperatures. However, the results have proven not to be qualitatively dependent on the specific choice of TPC model as long as rates are unimodal with exponential increases within the OTR.

### 4.3 Carbon Use Efficiency (CUE)

Species’ carbon use efficiency (CUE, henceforth denoted by *ε*) reflects the resource utilisation ability of a species. It is usually quantified as the partitioning of harvested resources into cell growth versus carbon loss mainly through maintenance (Geyer et al. 2016):

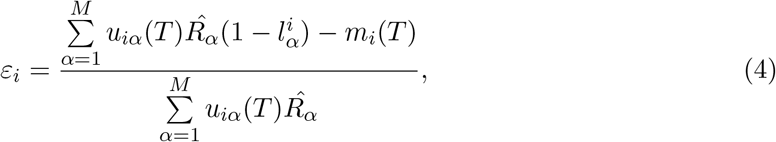

Where 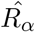 denotes the initial abundance of the *α*th resource and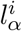here is a simplified representation of 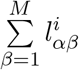 the total carbon or biomass lost during resource utilization due to the inefficiency of consumption as well as transformation of consumed resources into other resource types. Generally speaking, here, CUE is the ratio (proportion) of carbon allocated to growth accounting for loss through respiration and leakage (production of metabolic byproducts through the biochemical transformation of carbon resources) to that gained from uptake (the term in the denominator).

Here, metabolic traits (*u* and *m*) are temperature dependent and are denoted with (*T*). This means that the temperature performance of species’ CUE is solely determined by the thermal physiologies of metabolic traits.

### 4.4 Numeric Link between *R*^∗^ and CUE

According to the competitive exclusion principle, within a community where species exploit a certain resource niche, only species with relatively lower resource requirements could survive at equilibrium. In contrast, species with higher resource requirements for growth would be driven to extinction. This equilibrium resource requirement represents the relative fitness of the species within the community and is usually denoted as *R*^∗^. Here, we use a simplified quantification for *R*^∗^, which calculates each species’ intrinsic equilibrium resource requirement based on our microbial community model, considering each species growing on a single resource. The equation for our simplified *R*^∗^ standardized by the initial concentration of each resource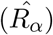 for species *i* is as follow:

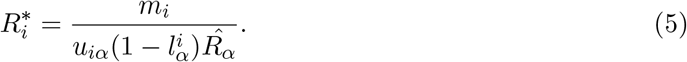

A simple rearrangement after combining equations 4 and 5 gives:

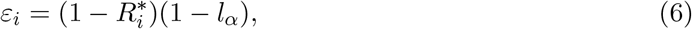

which suggests that species’ intrinsic CUE directly links to their relative fitness within the community (Figure 2 b). In this context, species’ metabolic strategies determine their competitiveness in species sorting during community assemblies under all given environmental conditions. This means that species with comparatively higher CUE also have lower equilibrium resource requirements (*R*^∗^) and therefore should survive in competitive exclusion.

### 4.5 Simulations

To investigate the effect of temperature on emergent species richness, we assembled heterotrophic bacterial communities under different temperatures through random immigration from a global species pool with different thermal metabolic strategies (TPCs of resources uptake rates and maintenance respiration rates). We simulate community assembly based on equation 1 and equation 2 with a random pool of 100 species competing for 50 carbon resources. For each assembly simulation, we numerically integrated the system until a dynamic equilibrium is reached for both the abundance of resources and the abundance of consumer biomass (which is *dC*_*i*_*/dt* = 0 and *dR*_*α*_*/dt* = 0), i.e., the surviving species coexisted.

In the MiCRM model, for each species, the uptake rates (*u*_*iα*_) of all resources are uniform random subsets of the species-level uptake rate (*u*_*i*_). This is achieved by setting *α* = 1 while assigning uptake preferences of resources using a Dirichlet distribution. This also allows all subset uptake rates of *M* resources to sum up to the species-level uptake rate. The leakage-transformation tensor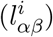is also sampled using a Dirichlet distribution, and the sum of the inefficiency of species-level resource utilisation for each resource 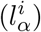 is 0.3. The sampling of both *u* and *l* is visually demonstrated in figure 1.

We parameterized the TPC of the species using data collected by Smith et al. (2019, 2021) from global online databases and empirical experiments in the laboratory. We used the *E* values for the species growth rates in these datasets as a proxy for the *E* values for the resource uptake rates. Empirical data on the thermal sensitivities of resource uptake rates are lacking; therefore, we assumed similar thermal sensitivities for uptake as those for growth rate. This is a reasonable assumption because the metabolic mechanisms of resource uptake by species mainly govern the maximum growth rates of microbial species. The parameterisation of TPCs of metabolic traits based on these datasets is explained in detail in Section SI.1 of the Supplementary Material. We particularly focused on the variance of normalisation constants, the variance of activation energies (standardised for uptake and maintenance respiration rates with the normalisation constants of both rates) and the covariance of these two parameters for both life-history traits (Figure SI.1).

The richness of emergent coexisting species at equilibrium is recorded at the end of each simulation as the number of species with biomass abundance *C*_*i*_ *>* 0. The uptake rates at species level and the maintenance respiration rate, CUE and *R*^∗^ of each species are collected alongside the emergent richness for each simulated assembly. The assembly simulations were replicated 670 times at each temperature ranging from 0 to 30 ^*o*^C.

All simulation code was written in the Julia language (Bezanson et al. 2017).

