## Supplementary Information for "The Role of Metabolic Strategies in Determining Microbial Community Diversity along Temperature Gradients"

### SI.1 Parameterization of the TPCs based on empirical datasets

We fit the data from Smith et al. (2019) using $T_r = 10^oC$ as the reference temperature for the TPCs of growth rates and maintenance processes, with which reference temperature we generated realistic TPCs for simulations. Based on this dataset, and the assumption that the thermal sensitivities of uptake are similar to those of the growth rate, the mean activation energy of the uptake rates ($E_u$) and the maintenance rates (respiration, $E_m$) are approximately 0.81 $eV$ and 0.57 $eV$. As uptake and respiration rates respond exponentially to temperature, the variation and covariance for their TPC normalisation constants are assumed to be log normal. The variation for both the normalisation constant and the activation energy of the uptake and respiration rates is standardised by their mean value. The coefficients for these variances ($\sigma(log(B_0))$ and $\sigma(E)$) are based on the growth rate data in this dataset, which are approximately 0.17 and 0.14. And the correlation coefficient ($\rho$) for the covariance between $log(B_0)$ and $E$ is approximately -0.35. The maximum performance temperatures $T_{pk}$ for the uptake and respiration rates are approximately 37.98 and 40.60 $^o$C in the data set; therefore, we parameterized the respiration rate TPCs to always be 3 $^o$C higher than the uptake rates.

However, this dataset does not contain enough data to calculate the normalisation constants of the uptake and respiration rates. Therefore, the mean log normalisation constant for the respiration rate $log(B_m)$ was collected from Smith et al. (2019), which is approximately -1.50. To calculate the mean normalisation constant for the uptake rate, we collected the mean CUE value in this dataset at our reference temperature ($\varepsilon_0$), which is approximately 0.22. Then with this $B_m$ and $\varepsilon_0$, and the calculation for CUE (equation for CUE see the Methods section) that $\varepsilon_0 = \frac{B_u(1-l)-B_m}{B_u} = 0.22$, we backcalculated the uptake rate at the reference temperature, which is the normalisation constant of the uptake rate $B_u$. The mean $log(B_u)$ is approximately -0.81.

### SI.2 Empirical Data and Randomly Generated Normalisation Constant and Activation Energy

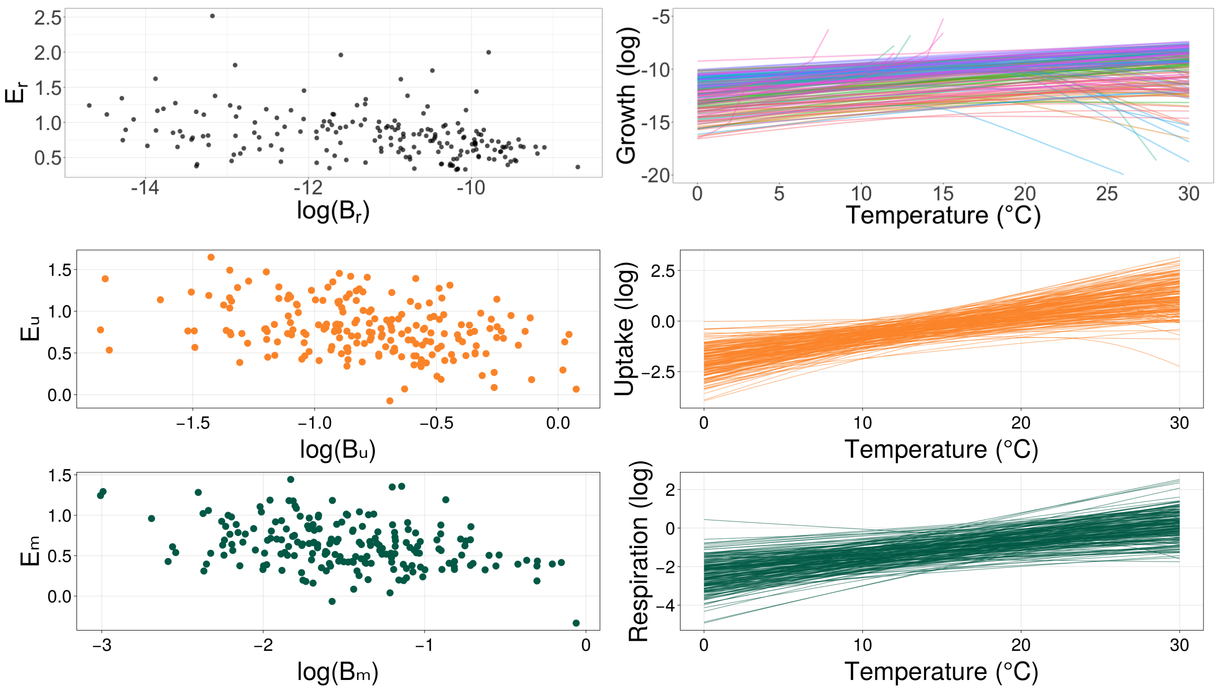


Figure SI.1: **Empirical metadata for $log(B_0)$ and $E$ of species growth rate and randomly generated $log(B_0)$ and $E$ parameter values for uptake and maintenance respiration rates.** The plot of both temperature dependency values is based on global metadata (black dots) on TPCs collected by Smith et al. (2019), which shows a slight negative covariance between $log(B_0)$ and $E$. The randomly sampled $log(B_0)$ and $E$ values for resource uptake rate and maintenance respiration rate based on the realistic variance and covariance analysed from the empirical datasets.

### SI.3 Shannon and Simpson Diversity Indexes

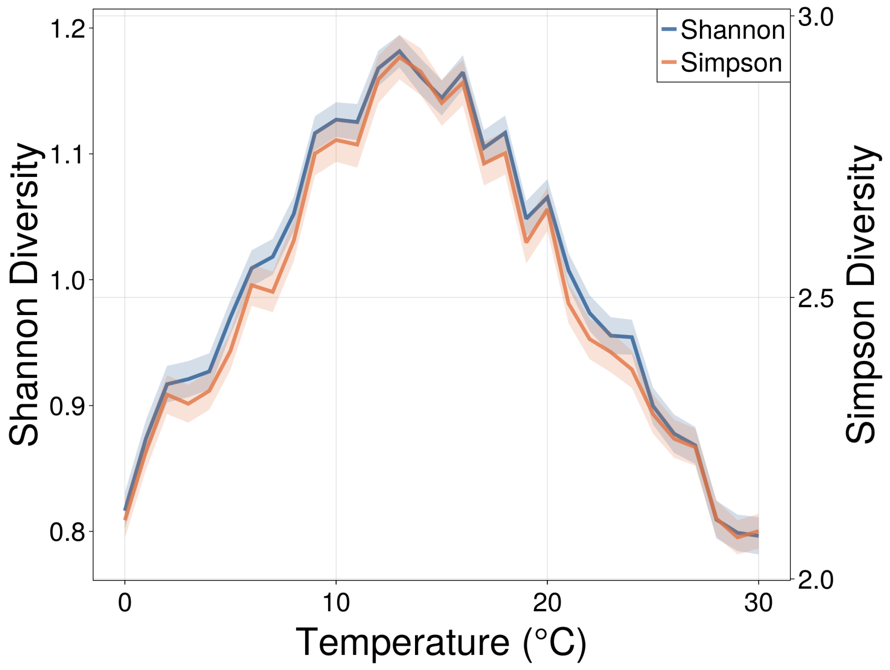


Figure SI.2: **Shannon and Simpson diversity along global temperature gradients.** Plotted with mean values taken from 670 simulations at each temperature with standard error.

### SI.4 Effective Lotka-Volterra and Feasibility

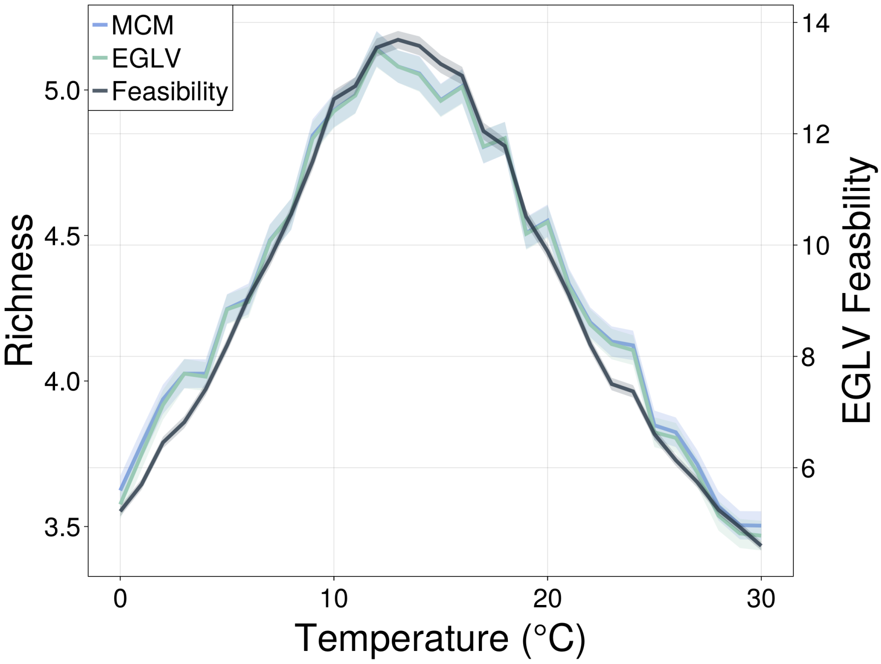


Figure SI.3: **Effective Lotka-Volterra simulation and the probability of feasibility.** The blue line depicts the mean richness predicted using the MCM, and the green line depicts the mean richness predicted using the effective Lotka-Volterra model with the original MCM parameterisation. The brown line depicts the calculated probability of feasibility using the mean-field approach (Clegg & Pawar 2022) on the effective Lotka-Volterra model. All lines are plotted with mean values taken from 670 simulations at each temperature with standard error.
